# Optimising CNT-FET Biosensor Design: Predictive Modelling of Biomolecular Electrostatic Gating and its Application to Beta-Lactamase Detection

**DOI:** 10.1101/2023.07.31.551308

**Authors:** Rebecca E.A. Gwyther, Sébastien Côté, Chang-Seuk Lee, Krithika Ramakrishnan, Matteo Palma, D. Dafydd Jones

**Author notes:** These authors contributed equally (Rebecca E.A. Gwyther, Sébastien Côté, Chang-Seuk Lee). Corresponding authors: DDJ: Molecular Biosciences Division, School of Biosciences Cardiff University, Cardiff, CF10 3AX, UK., MP: Department of Chemistry, School of Physical and Chemical Sciences Queen Mary University of London, UK., SC: Département de Physique, Cégep de Saint-Jérôme, Saint-Jérôme (QC), J7Z 4V2, Canada.

## Abstract

Carbon nanotube field effect transistor (CNT-FET) setups hold great promise for constructing next generation miniaturised biosensors whereby a biomolecular event gates conductance. The main issue is predicting how proteins, with their innate mosaic and distinctive electrostatic surfaces, interact with and thus modulate conductance of the CNT-FET. To overcome this barrier, we used advanced sampling molecular dynamics combined with non-canonical amino acid chemistry, to model the protein electrostatic potential imparted on SWCNTs. Here, we focused our efforts using β-lactamase binding protein (BLIP2) as the receptor due to its potential as a biosensor for the most common antibiotic degrading enzymes, the β-lactamases (BLs). Modelling was confirmed experimentally by attaching BLIP2 at single designed residues positions directly to SWCNTs using genetically encoded phenyl azide photochemistry. Our devices were able to successfully detect the two different BLs, TEM-1 and KPC-2, with each BL generating distinct conductance profiles due to differences in their unique surface electrostatic profiles presented close to the SWCNT surface. The changes in conductance closely matched the predicted electrostatic profile sampled by the SWCNTs on BL binding. Thus, our modelling approach combined with new and straight-forward residue-specific receptor attachment techniques, provides a general approach for more effective and optimal CNT-FET biosensor construction.

## Introduction

Nanocarbon devices such as carbon nanotube field effect transistors (CNT-FETs) are rapidly becoming the base biosensing material of choice due to their desirable electronic properties and routes to functionalisation ^1–4^ enabling label free detection of biomolecules with high sensitivity to specific targets using miniaturised devices ^5–10^. In a CNT-FET biosensor setup, biomolecular interaction events are detected due to the electrostatic field generated by the biomolecules in proximity of the CNT’s surface inducing changes to charge carrier density and thus modulating CNT conductance. ^2, 6, 11^ To enable target specificity, single walled CNTs (SWCNTs) are typically decorated with receptors (e.g. DNA aptamers ^12–16^ or binding proteins ^17–21^) that recognise and bind the required analyte. In this regard, protein-protein interactions (PPIs) are of particular interest as they are widespread in nature being central to a range of biological events and are at the heart of modern diagnostics.

Proteins have surfaces comprised of basic (positive) and acidic (negative) residues giving rise to an electrostatic surface unique to each protein. Whilst there have been notable successes of proteins electrostatically gating SWCNTs ^17–19^, their effectiveness is generally limited by how the receptor is attached to the SWCNT surface. Ideally, the receptor should be attached in a defined manner (i.e. at a predefined site), with an optimal orientation to impart the greatest electrostatic effect onto the SWCNT upon analyte binding. However, most receptor protein attachment is not controlled and is essentially random leading to heterogenous orientations of the receptor (and in turn the analyte), of which many will be far from optimal (see Figure 1a for examples), generating signal variation from device to device, and even signal cancelling effects. There have been some notable successes regarding designed attachment via specified residues, usually cysteine thiol chemistry ^18, 19^ or more recently non-canonical amino acid chemistry incorporated using a reprogrammed genetic code approach ^17, 22, 23^. The latter is particularly powerful choice as there is a broader selection of coupling chemistry available and can also allow attachment either directly to the SWCNT ^22, 23^ or via an intermediate linker ^17, 24, 25^. For example, genetically encoded phenyl azide chemistry ^26^ enables direct photochemical covalent attachment ^23^ or linker-based click chemistry ^17, 25^ approaches to be used.

**Figure 1.**
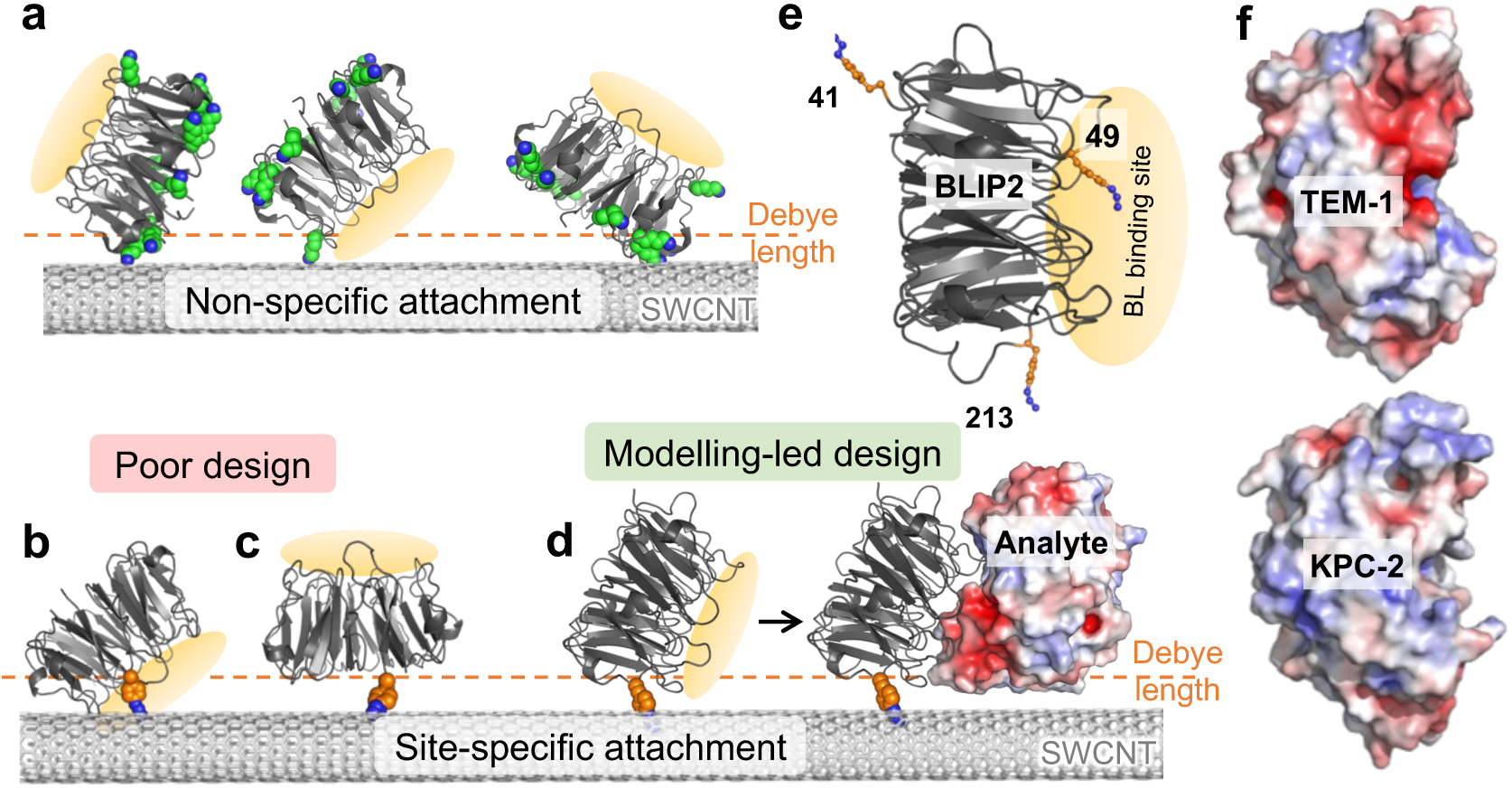
Outline of the CNT-FET design. (a) Examples of non-specific receptor protein (BLIP2) attachment via primary amines with lysine residues shown as green spheres. The Debye length is shown as a dashed orange line. (b-c) Examples of poor design for site-specific attachment whereby the BLIP2 binding interface is blocked by the SWCNT (b) or binding of the analyte will be well beyond the Debye length (c). (d) Example of an ideal, modelling-led design whereby the analyte binding is within the Debye distance and imparts a predicted electrostatic profile on the SWCNT. (e) Structure of BLIP2 highlighting the BL binding interface (orange oval) and selected interface residues (orange sticks) mutated to azF. (f) Different surface electrostatic profile of two BLs, TEM-1 (top) and KPC-2 (bottom). The equivalent surfaces are shown for each. Red and blue represent acidic and basic patches, respectively.

The main issue with residue-specific attachment is picking the right residue to maximise biosensing potential so as to avoid: (1) steric clashes between receptor protein and/or analyte protein and the SWCNT; (2) no analyte binding due to masking of the receptor binding site (Figure 1b); (3) analyte binding far beyond the Debye distance (Figure 1c). The Debye length is particularly important as it defines the distance from the SWCNT where the electrostatic gating is greatest ^16, 17, 27^. Thus, modelling how a receptor protein is attached and how the subsequent analyte binds with respect to the SWCNT is critical to achieving optimal CNT-FET designs (Figure 1d).

Herein we focused our efforts on the detection of a group of enzymes, β-lactamases (BLs), that are one of the main causes of antimicrobial resistance (AMR) ^28–30^ Class A β-lactamases, such as the clinically important TEM-1 ^31^ and KPC-2 ^32^, are the main causes of resistance to β-lactam antibiotics so their quick detection during infection could prove vital in administering the right antibiotic therapy. The β-lactamase inhibitory protein 2 (BLIP2; Figure 1e) ^33, 34^ is a near “universal” binder for class A BLs so acts as the ideal receptor protein for detecting a broad range of BLs, including TEM-1 ^35^ and KPC-2 ^36^. While TEM-1 and KPC-2 are structurally and functionally very similar, they have distinctive electrostatic surface profiles (Figure 1f).

By combining computational modelling approaches, genetic code reprogramming, chemical biology, and nanoscale fabrication of biosensing devices, we successfully photochemically attached BLIP2 at defined, designed residues, enabling the detection of TEM-1 and KPC-2; each BL generated distinct conductance profiles due to differences in their unique surface electrostatic patches presented close to the SWCNT surface; the changes in conductance closely matched the predicted electrostatic model profiles sampled by the SWCNTs on BL binding.

## Results and discussion

### Modelling receptor protein attachment site feasibility

We have recently developed an approach whereby a protein can be precisely and intimately interfaced directly with nano-carbon, including SWCNTs, using genetically encoded phenyl azide chemistry via the non-canonical amino acid azF (4-azido-L-phenylalanine) ^22, 23, 37^. Here we aimed to devise an accurate method to model the likely initial binding orientation of the receptor protein allowing us to estimate the distance and thus electrostatic influence between the incoming analyte protein and the SWCNT. BLIP2 will act as the receptor protein on the CNTs, for both its potential in antimicrobial resistance (AMR) diagnostics, and to test our modelling approach. As BLIP2 binds a range of different BLs, this allowed us to model how different analyte proteins with distinctive electrostatic surfaces influence SWCNTs and thus conductance.

The first step is to assess whether a particular attachment residue on the receptor protein [BLIP2] is viable for interfacing with a SWCNT by using molecular dynamics to sample rotamer forms of azF. We then used a more extensive electrostatic profiling sampled by the SWCNT on analyte protein [the BLs] binding to predict the electrical response of the CNT-FET. Structures available in the Protein Data Bank (PDB) (e.g. BLIP2-TEM-1 complex ^35^) or models generated by in silico methods such as AlphaFold2 (AF2) multimer ^38–40^ (e.g. BLIP2-KPC-2; Figure S1) can be used as the starting point. The selected BLIP2 residues are then mutated in silico to incorporate azF and the BLIP2 receptor variant structures are used as starting points for molecular dynamics to sample the rotamer configurations of the azF side-chain. We selected two BLIP2 residues, 41 and 213 (Figure 1b), as models to test our approach. We have shown previously that site-specific attachment via these residues through an intermediary pyrene linker allowed detection of TEM-1 in a CNT-FET setup. ^17^

MD simulations suggested that BLIP2 with azF at residue 41 (BLIP2^41azF^) sampled two major rotamer conformations covering 92.5% the sampled frames (Figure 2a-b). Docking of rotamer 1 (R1) onto a SWCNT suggested significant steric clashes between the protein and nanotube (Figure S2). No steric clashes were observed when rotamer 2 (R2) was manually docked to the SWCNT. While R1 was the less populated of the two major rotamer forms, it is sampled on a regular basis during the simulation (Figure 2a). Docking of TEM-1 onto R2 BLIP2^41azF^-CNT suggested that the analyte BL would still bind to the receptor-CNT complex (Figure 2c).

**Figure 2.**
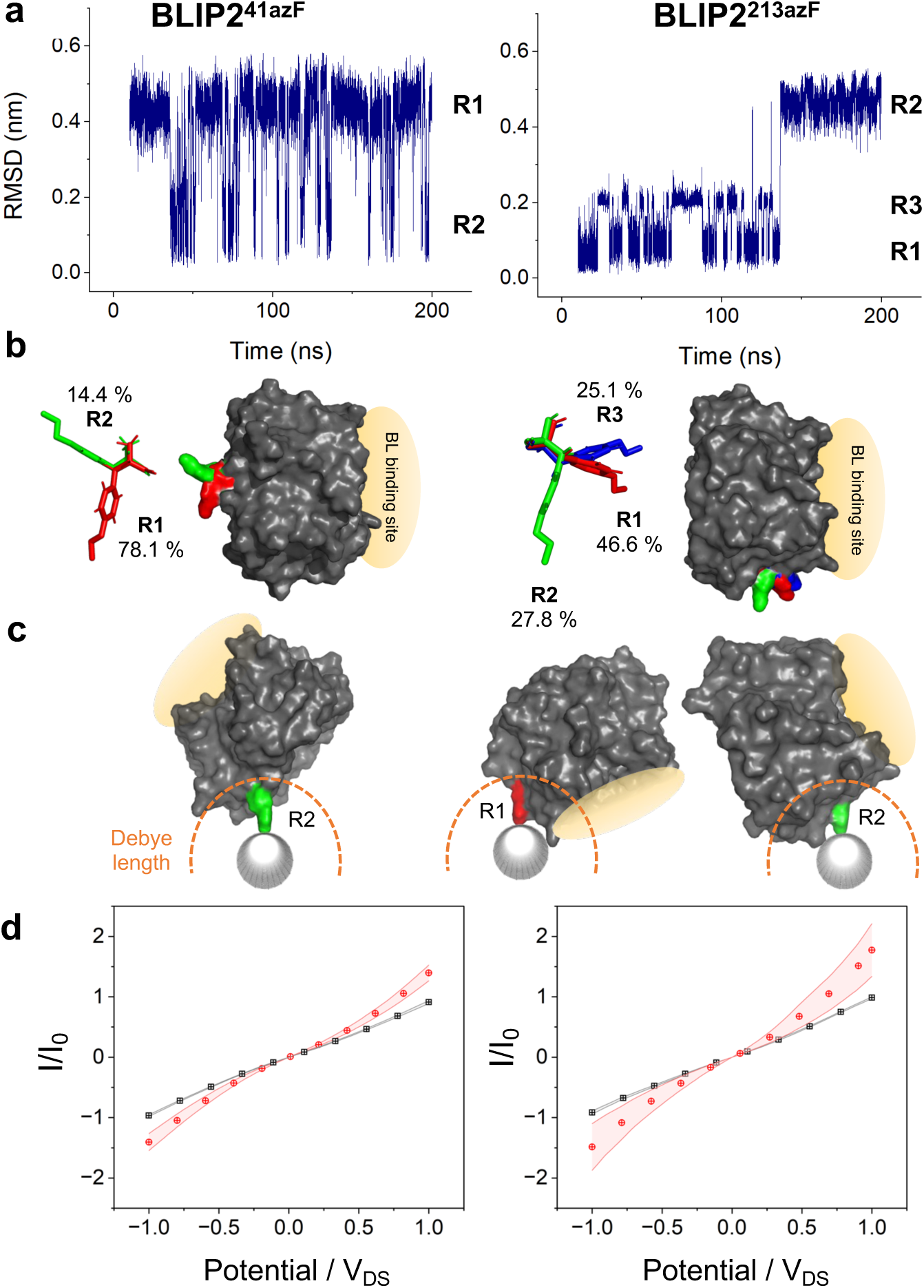
Modelling of BLIP2 mutants (azF41 left column and azF213 right column) azF sidechain rotamer configurations and SWCNT binding. (a) Root mean squared deviation (RMSD) for the azF residue over one simulation repeat (190 ns) in BLIP2^41azF^ and BLIP2^213azF^, with rotamer states identified from step changes in RMSD. (b) Modal rotamer configurations were extracted and compared across all repeats (n=3) to give an average rotamer propensity, labelled as percentage of total simulation time. (c) Viable rotamers for SWCNT docking. Docking of BLIP2 azF mutant onto the SWCNT is based on the geometry of the known [2+1] cycloaddition reaction, whereby the nitrene radical formed on exposure to UV light inserts perpendicular to the SWCNT ^41^. Rotamers were deemed viable if no steric clash occurred between BLIP2 and the SWCNT. The BL binding site is highlighted in yellow and Debye length is depicted in orange. (d) I/Vs before (black) and after (red) BLIP2 azF attachment; BLIP2^41AzF^ (n=15) and BLIP2^213azF^ (n=8). Standard error of the mean is plotted either side of the average measurements.

BLIP2 with azF at residue 213 (BLIP2^213azF^) accessed 3 main rotamer conformations (Figure 2a-b) accounting for 99.5% of the frames. Only rotamer 3 (R3) displayed steric clashes when manually docked on the SWCNT (Figure S2). On docking BLIP2^213azF^ R1 and R2 forms to the SWCNT, both rotamer forms retained the ability to bind TEM-1. This initial modelling suggested that R1 brought the BL close to the SWCNT surface while R2 caused the BL binding site of BLIP2^213azF^ to project away from the nanotube (Figure 2c).

We then experimentally confirmed that both BLIP2^41azF^ and BLIP2^213azF^ could be photochemically attached to SWCNT bundles in the CNT-FET setup. We have previously shown that BLIP2^41azF^ could be attached to single SWCNTs by AFM ^23^, which we confirmed here with SWCNT bundles for both BLIP2^41azF^ and BLIP2^213azF^ (Figure S3 and associated discussion). Additional height increases were observed on the addition of TEM-1 to the same SWCNT bundles (Figure S3) providing evidence that BL binding capacity is retained as the initial modelling predicted. Photochemical BLIP2 azF attachment was also confirmed by I/V measurements in a CNT-FET setup using p-type semiconducting SWCNTs (Figure 2d). The I-V measurements pre-functionalisation display ohmic behaviour; with a linear relationship between current and voltage, as expected with pristine SWCNTs. Covalent attachment of either BLIP2 azF variants resulted in increased conductance as expected given that both attachment sites are predicted to impose a negative electrostatic potential on the SWCNT (vide infra). The introduction of breaks into the SWCNT sp^2^ bond network would normally lead to a reduction in conductance; by using the phenyl azide photochemical approach, our work suggests that this can be offset, with recent observations indicating aromatic azides maintain the π electron network upon covalent functionalisation ^41^.

## Modelling electrostatic effect of protein on SWCNT

Predicting the electrostatic influence proteins have on the SWCNT conductance channel is critical to the success of FET-based biosensors. Here we have developed an approach to model electrostatic effects of protein binding to a SWCNT. The approach used: (1) exhaustively sampling the orientations of receptor protein with respect to the carbon nanotube using all-atom, solvent explicit Hamiltonian-replica exchange molecular dynamics (H-REMD) simulations^42^ and; (2) determining the electrostatic potential generated by these orientations on the surface of the SWCNT using electrostatic Poisson-Boltzmann simulations^43^. Using this procedure, we quantified the electrostatic change due to attachment of BLIP2 via azF41 or azF213 to the SWCNT and the subsequent association of TEM-1 and KPC-2. From the electrostatic potential, we then infer the impact on the charge carrier density in the SWCNT and thus on its conductance.

First, the 750 ns per replica H-REMD simulations thoroughly sampled a variety of orientations that progressively converged towards a few thermodynamically stable clusters over the last 250 ns: BLIP2^41azF^ and BLIP2^213azF^ have respectively two and four main orientation ensembles (Figure S4). Inspection of the centroid of these ensembles reveals that TEM-1 is localized very differently with respect to the nanotube when associating to BLIP2^41azF^ compared to BLIP2^213azF^ (Figures S5 and S6). Indeed, while all residues of TEM-1 are farther than 1.5 nm from the nanotube for BLIP2^41azF^, many charged residues are within 1.5 nm of the nanotube for BLIP2^213azF^. Consequently, we expect that the electrostatic potential change on the nanotube will be significantly stronger when the analyte (TEM-1) binds to the BLIP2^213azF^ as the receptor.

Next, we quantified the electrostatic potential (ESP) change by solving the Poisson-Boltzmann equation for a total of 2500 representative orientations from the H-REMD simulation for each attachment site. Binding of either BLIP2^41azF^ or BLIP2^213azF^ alone imparts a largely negative ESP on the SWCNT (Figure S7), in line with increased conductance of p-type SWCNTs observed in Figure 2d. Binding of TEM-1 elicits an average reduction of the ESP on the order of 1 mV on the nanotube for BLIP2^41azF^, while the reduction is an order of magnitude larger for BLIP2^213azF^ (10 mV) (Figure 3a). To quantitatively compare the two sites, we calculated the global ESP change on the surface of the nanotube (Figure 3b). Binding of TEM-1 produces a relatively small negative global ESP change with a peak at –2,700 ± 420 a.u for BLIP2^41azF^; this is significantly stronger for BLIP2^213azF^ with two peaks at –8,649 ± 1,991 a.u and –28,020 ± 4,848 a.u.

**Figure 3.**
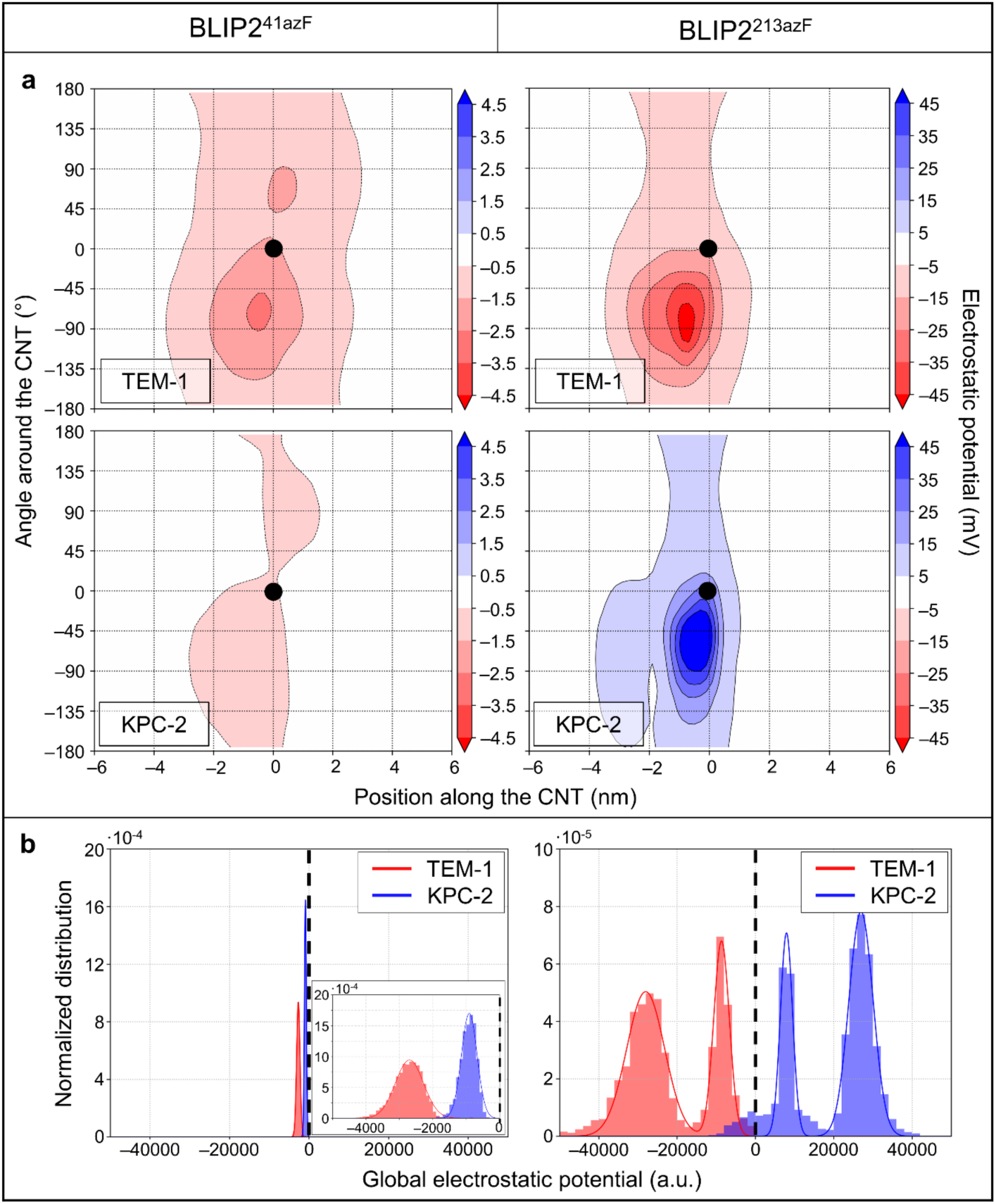
Change of electrostatic potential on analyte binding. The change of electrostatic potential (ESP) on the carbon nanotube due to the association of TEM-1 and KPC-2 with BLIP2^41azF^ (left) and BLIP2^213azF^ (right). The BLIP2 alone ESP profiles are shown in Figure S7. (a) The average electrostatic potential exerted by each BL as a function of the position along the nanotube axis and of the angle around the nanotube where the coordinate (0,0) corresponds to the attachment site (black dot). The average is performed on the electrostatic potential maps of 2500 sampled configurations in the converged interval (see SI). (b) The normalized histogram of the global ESP of each BL on the nanotube. The global electrostatic potential is computed using a Riemann sum for each sampled configuration separately. The black dashed line indicates no ESP change compared to BLIP2 alone. For BLIP2^41azF^ binding a BL, inset is the magnified region of the global electrostatic potential.

As shown in Figure 1f, KPC-1 has a distinct electrostatic surface profile to TEM-1, with KPC-2 having an overall charge of –1, while TEM-1 is –7. Our simulations indicate that this difference in surface charge profile for KPC-2 has a strong impact on the ESP imparted on the nanotube, particularly for BLIP^213azF^ for which the ESP profile becomes positive instead of being negative (Figure 3a). In term of overall ESP, two peaks are again observed for KPC-2 on binding BLIP2^213azF^ but now with a significant shift to a positive ESP sampled by the SWCNT (26,971 +/-3,095 and 7945 +/-1558 a.u.; Figure 3b); association of KPC-2 results in a relatively small negative overall ESP on binding BLIP2^41azF^ (–908 +/-234 a.u.).

Thus, our modelling predicts that TEM-1 binding to BLIP2^213azF^ is likely to exert the greatest ESP on SWCNT; the increased negative charge close to the p-type SWCNT conductance channel should increase the positive hole carrier density and so increase conductance ^16, 17, 44, 45^. TEM-1 binding to BLIP2^41azF^ should only exert a small negative ESP so should generate a relatively small increase in conductance. In comparison, KPC-2 binding to BLIP2^213azF^ will impart a significant positive charge on the p-type SWCNT so reducing conductance. Binding of KPC-2 to BLIP2^41azF^ should exert a small negative ESP which will potentially have a very minor effect on conductance.

## Conductance of designed BLIP2 receptor CNT-FETs

We next tested our models to see if conductance changed in the anticipated manner in a CNT-FET setup. All CNT-FETs used were comprised of p-type SWCNTs, and I-V traces were measured on addition of the receptor [BLIP2; Figure 2d] and then analyte [BL] (Figure 4a).

**Figure 4.**
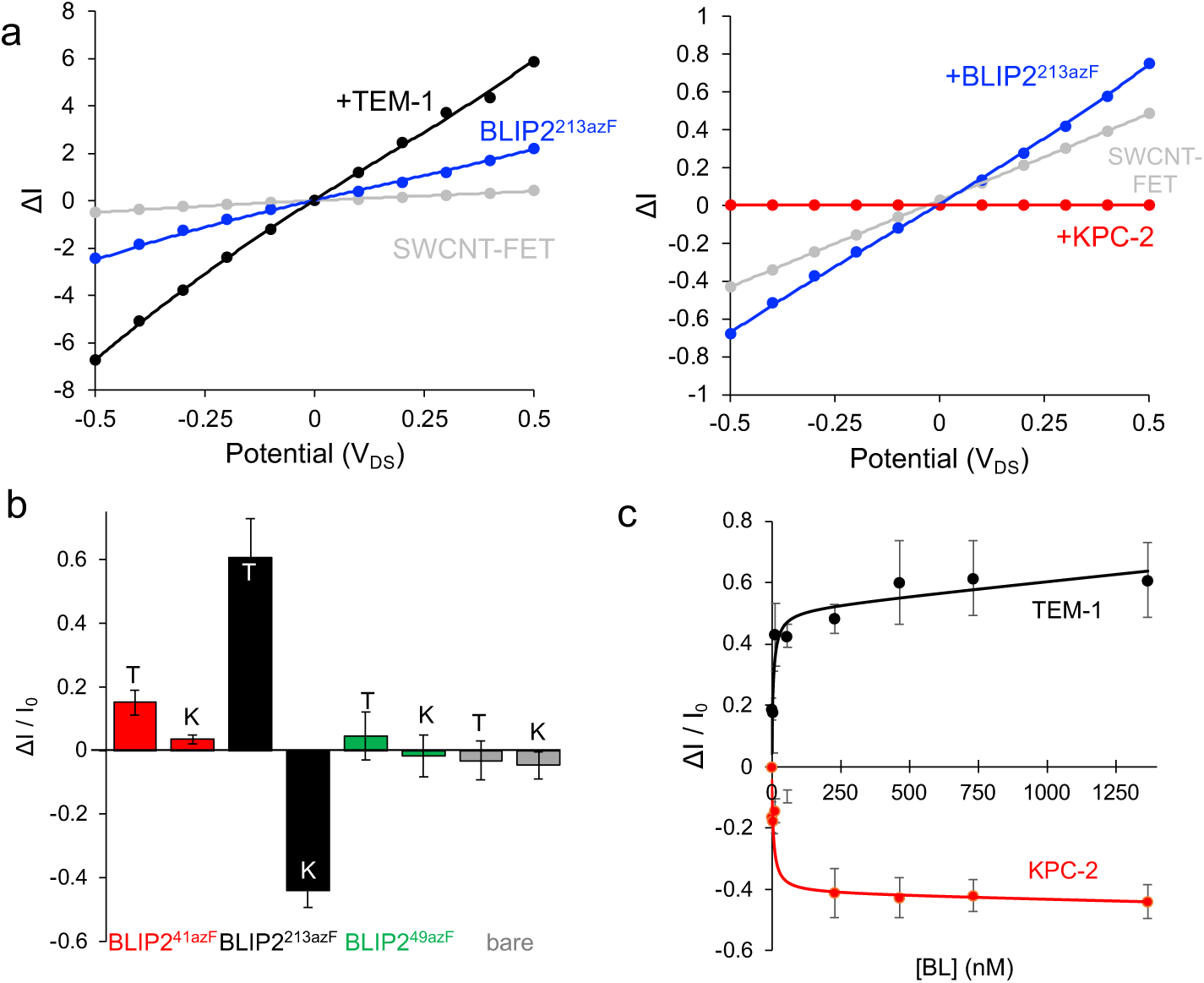
Conductance measurement with BLIP2 azF variants photochemically attached to CNT-FET devices upon the addition of β-lactamase. (a) Example I-V traces for BLIP2^213azF^ binding TEM-1 (left) and KPC-2 (right). The red (squares), blue (circles) and orange (triangles) plots represent bare CNT-FET, attachment of BLIP2^213azF^ and addition of 1300 nM TEM-1, respectively. (b) Relative changes in conductance on addition of either TEM-1 (T) or KPC-2 (K) to either bare CNT-FETs (grey) or with BLIP2^41azF^ (red), BLIP2^213azF^ (black), BLIP2^49azF^ (green) acting as the SWCNT bound receptor. Each BL was added to a final concentration of 1.4 μM. Error bars are standard deviations between independent readings on 3 different devices. (c) BL concentration-dependent change in current (at V_DS_ of 0.1 V) plots for BLIP2^213azF^. TEM-1 and KPC-2 conductance profiles are coloured black and red, respectively. The conductance data were collected via I-V measurements between source-drain. The data points where fit to one site binding equation in GraphPad Prism. Error bars are standard deviations between independent reading on 3 different devices at each concentration.

Notably, on addition of a BL, the I-V traces for BLIP2^213azF^ devices gave the largest changes in conductance compared to devices with BLIP^41azF^ (Figure 4b), in line with the ESP models. Additionally, the BLIP2^213azF^ devices behaved differently depending on the BL added. Addition of TEM-1 resulted in increased conductance whereas KPC-2 caused conductance to decrease (Figure 4b). This matches our predictions whereby TEM-1 binding to BLIP2^213azF^ exerts a negative ESP on the SWCNT (Figure 3b) so enhancing positive hole charge carrier mobility our p-type devices resulting in increased conductance. KPC-2 on the other hand exerts a positive ESP on the SWCNTs which will decrease conductance.

TEM-1 binding to BLIP2^41azF^ resulted in only a small conductance increase (Figure 4b and Figure S8), 4-fold lower than that observed for BLIP2^213azF^ devices. Modelling suggests that TEM-1 binding to BLIP2^41azF^ exerts a relatively weak negative ESP profile on the SWCNT, so our experimental results are consistent with the models (Figure 3b). KPC-2 exerts an even weaker ESP on SWCNTs, and this would explain the negligible change conductance (Figure 4b and Figure S8).

We also used a BLIP2 variant, BLIP2^49azF^, as a control receptor that is known to sterically block BL binding on SWCNT attachment^17^; the azF photochemical attachment handle is within the BL binding region (Figure 1e). As shown in Figure 4b (and Figure S9), little change in conductance is observed on addition of either BL to BLIP2^49azF^ CNT-FETs, as expected. Furthermore, each BL does not induce significant changes in conductance of bare SWCNTs highlighting the importance of the receptor (Figure 4b and Figure S9).

We next investigated BL concentration dependent changes in conductance for each device setup. Initially Dulbecco’s phosphate buffered saline (DPBS) buffer was added to the channel region followed by stepwise additions of the BLs, with I-V curves measured after every BL addition (Figures 4c, S8 and S9). For the BLIP2^213azF^ receptor devices, we observed a TEM-1 concentration dependent change in conductance (Figure 4c). Even at low TEM-1 concentrations (sub 100 nM) close to maximal signal was observed suggesting saturation of BLIP2 binding sites and retention of high affinity binding observed for native BLIP2 to BLs. ^46, 47^ As noted above, we did see a smaller increase in conductance for TEM-1 binding BLIP2^41azF^ which did show a concentration dependence (Figure S8). For KPC-2, a similar concentration dependence was observed with BLIP2^213azF^ CNT-FETs, but with a decrease in conductance now used to monitor BL binding (Figure 4c). Again, signal saturation was reached at relatively low concentrations of KPC-2 suggesting that BLIP2^213azF^’s high BL affinity was retained. No clear KPC-2 binding signal was observed with BLIP2^41azF^ modified CNT-FETs (Figure S8) suggesting there was no ESP change on the SWCNT. Both bare SWCNTs and BLIP2^49azF^ also showed no BL concentration-dependent signal change (Figure S9).

## Conclusions

While the use of a bioCNT-FET setup offers great potential for biosensing and even in molecular electronics, it is of paramount importance to design and predict device performance. The standard route of non-specific attachment of protein receptors leads to a highly heterogenous system in terms of receptor orientation, and thus analyte binding that may even result in electrostatic potential effects cancelling each other out, so impairing the optimal performance of the biosensor. Here we show that in silico modelling has the potential to predict accessible receptor attachment sites and the likely ESP sampled by a SWCNT – acting as the transducer in FET biosensors that manifests as expected conductance changes. Critical is the use of a non-natural amino acid that restricts the coupling site to a single defined residue on the receptor providing a homogenous and precise link to the SWCNT, and so defining the modelling process. Moreover, we show that similar proteins but with differing surface ESP profiles elicit discrete conductance profiles, an aspect that we can also predict through modelling. Here we show that two different BLs with differing clinical effects in terms of their resistance profile generated different conductance profiles. This could pave the way for rapid sensing of BLs based on their electrostatic profiles present during infection leading to more appropriate antibiotic prescription regimes. We envisage that our combined modelling and non-natural amino acid approach could be applied more broadly to model the likely spatial and functional relationship of protein-bionanohybrid systems so helping to understand fundamental basis of action and generating more effective designs for better performing bioCNT-FETs.

## Methods

### Modelling BLIP2 azF mutations and rotamers

Input structures were prepared using AlphaFold v2.1.0 ^39^ using the translated gene sequence of BLIP2, to most accurately represent the protein recombinantly produced. The structures produced are highly similar to the known crystal structures of BLIP2 with an all atom RMSD of 0.461 and 0.455 Å for BLIP2^41azF^ and BLIP^213azF^, respectively, compared to PDB entry 3qi0 ^47^.The azF mutation was introduced manually using the SwissSidechain plugin in PyMOL ^48^. All simulations were performed in GROMACS ^49, 50^ with the CHARMM36 forcefield ^51^. The MD simulations were performed as outlined in the SI. Post-MD, BLIP2 was centred in the box as treatment for PBC and fitted to the reference file by translation. Structural stability analysis was performed to test the viability of the model, and the first 10 ns were trimmed from all trajectories to discount the equilibration phase of the simulation.

### H-REMD simulations

All-atom, solvent explicit Hamiltonian replica-exchange molecular dynamics (H-REMD) simulations were performed on BLIP2 covalently attached to a carbon nanotube for the two attachment sites (41azF and 213azF). Below, we briefly present the simulated systems with more detailed descriptions in the SI.

The protein is modelled using the AMBER14sb force field ^52^. The attachment between BLIP2 and the nanotube is parametrized using the GAFF/RESP procedure from AmberTools20 ^53^. The nanotube atoms are modelled as uncharged sp2 carbon of type CA in AMBER, with Lennard-Jones parameters σ=0.339967 nm and ε=0.359824 kJ/mol, and their position is frozen during the simulation. Water molecules and ions are respectively modelled using TIP3P and Joung-Cheatham parameters.^54^

The H-REMD simulations were performed using the GROMACS software ^50, 55^ version 2019.6 augmented with the open-source, community-developed PLUMED library ^56, 57^ version 2.6.2 for H-REMD simulations ^58^. The simulations were done in the NVT ensemble using the Bussi-Donadio-Parrinello thermostat ^59^ with a reference temperature of 300 K. Bonds involving a hydrogen atom were constrained using the P-LINCS algorithm and the geometry of the water molecules was constrained using the SETTLE algorithm, allowing an integration timestep of 2 fs. The cutoffs for the short-range Lennard-Jones and electrostatic interactions were 1.0 nm. Long-range electrostatic interactions were computed using the smooth particle mesh Ewald method ^60^.

One H-REMD simulation of 18 µs (0.75 µs per scale, 24 scales) was performed for each covalent BLIP2 attachment site (azF41 and azF213) to the nanotube. To achieve thorough sampling, the energy scales progressively reduce the attractive part of the Lennard-Jones interactions between BLIP2 and the carbon nanotube. Consequently, this improves the dissociation of BLIP2 from the nanotube at larger scaling, allowing it to sample an ensemble of orientations with respect to the nanotube as the system diffuses back to lower scaling, while preserving the overall structure of BLIP2. Specifically, there are 24 scales ranging from 1 (no scaling, original energy) to 0 (no Lennard-Jones attraction between BLIP2 and the nanotube): 1.00, 0.95, 0.91, 0.87, 0.84, 0.81, 0.79, 0.76, 0.73, 0.71, 0.68, 0.66, 0.63, 0.60, 0.55, 0.50, 0.45, 0.40, 0.34, 0.28, 0.22, 0.15, 0.08 and 0. The scales were optimized to yield an average exchange rate of approximately 30% between scales 1 and 0.87, 55% between scales 0.87 and 0.60, and 30% between scales 0.60 to 0. The intermediate scales ensure a smooth transition from BLIP2 having many interactions with the nanotube (scales 1 to 0.87) to BLIP2 being mostly detached from the nanotube (scales 0.60 to 0). Exchanges between neighbouring scales are tried each 10 ps, alternating between even-odd and odd-even scales. The initial structures used for the scales are presented in Figure S10 (SI).

### Electrostatic potential simulations

The APBS software was used to solve the non-linear Poisson-Boltzmann equation ^43^ to determine the electrostatic potential generated by the protein on the SWCNT. The structures sampled at the unscaled energy scale using H-REMD were taken with a timestep of 100 ps in the 500-750 ns interval (2500 structures in total for each simulation) for electrostatic analysis. The solvent was removed, keeping only BLIP2, the attachment and the SWCNT. To compute the effect of TEM-1 on the electrostatic potential generated on the nanotube, TEM-1 was added by aligning the structure of the BLIP2/TEM-1 complex onto the structure of BLIP2 in the frames sampled during the simulations. The aligned TEM-1 was kept along with the simulated BLIP2. For KPC-2, the same was done to obtain the BLIP2/KPC-2 complex as it shares high structural similarity with the BLIP2/TEM-1 complex according to AlphaFold predictions (Figure S1). The electrostatic potential calculations for BLIP2, BLIP2/TEM-1 and BLIP2/KPC-2 use the same parameters and box size (see SI for more details).

### Protein engineering and purification

The BLIP2 AzF variants were produced and purified as described previously ^17, 23^. It is important to note that the production and purification of BLIP2 azF variants was performed under dark conditions where possible to minimise light exposure, and UV light sources were turned off during chromatography. TEM-1 was expressed using a pET-24a vector (a kind gift from the Makinen lab)^61^ in *E. coli* BL21(DE3) cultured in 2xTY media supplemented with kanamycin (30 mg/ml). On reaching an OD600 of ∼0.6, 1 mM IPTG was added to induce expression – kanamycin. The culture was then incubated at 22°C for 16 hours. The next day, cells were pelleted at 5000 rpm and 5°C for 20 minutes in the Fiberlite^TM^ F9-6 x 1000 LEX rotor, and the supernatant removed. The pellet then underwent periplasmic extraction to carefully extract the protein, without lysing the whole cells as described in the SI. A similar procedure was used for KPC-2 except chloramphenicol (12.5 μg/ml) was used during cell culture. The BL proteins were then purified by a combination of ion-exchange chromatography and size exclusion chromatography as outlined in the SI.

### Device fabrication and protein attachment

Electrodes were patterned as previously reported ^17, 62^ and purchased from ConScience. Nanosized electrodes were fabricated on a p-doped Si/SiO_2_ wafer by a combination of laser and electron beam lithography, followed by evaporating a thin adhesive layer of Cr and a thick layer of Au. 0.1 mg of 95% semiconducting single walled carbon nanotubes (SWCNTs) was dispersed in 1 mL of 1% SDS solution via sonication for 1h. The SWCNT sample was centrifuged for 1 h and the supernatant was collected and used as stock solution. To immobilise a small bundle of SWCNTs between electrodes, dielectrophoresis (DEP) was performed by applying an alternating current (AC) voltage between electrodes after SDS-dispersed SWCNT solution was cast on the electrodes. Typically, the frequency of the generator was switched onto Vp-p=3V at f=400kHz. After DEP, immobilised SWCNTs were characterised by AFM measurements. Protein attachment was performed by applying UV light after a 1 μM BLIP2 protein solution was cast on the CNT-FETs. After 5 min of UV light exposure, the substrate was washed with water for 5 min to remove excess proteins from the substrate and blown gently with nitrogen gas. Topography analysis of the electrodes were imaged with a Bruker Dimension Icon atomic force microscope (AFM) with ScanAsyst Air tips. The images were analysed with NanoScope Analysis.

### Conductance measurements

Conductance measurements were obtained with a probe station (PS-100, Lakeshore) equipped with a semiconducting parameter analyser (Keithley, 4200 SCS) at room temperature. I-V curves were measured with bias sweeping mode (–1 V to 1 V) applied to source electrode while drain electrode was grounded. For real-time measurements, 100 mV source-drain bias was applied across the devices which have been covered with a drop of DPBS buffer solution. DPBS buffer was first added to the devices after reading of the current was stable, which cannot induce changes in current. This stable reading of the current through devices was recorded as background. Subsequently, 5µL of TEM-1 and KPC-2 solutions of various concentrations were cast on the devices. The resulting conductance profiles were collected by calculating ΔI/I_0_ (ΔI/I_0_ = (I_meas_ – I_0_) / I_0_, where I_meas_ is the measured conductance after the addition of the target analyte, and I_0_ is the measured conductance with only DPBS buffer solution added. The data was then fitted to a one-site total binding equation in GraphPad Prism.

## Acknowledgements.

R.E.A.G., S.C. and C-S.L. contributed equally to this work. D.D.J would like to thank the EPSRC (EP/J015318/1 and EP/V048147/1) for supporting this work. We gratefully acknowledge financial support from the U.S. Air Force Office of Scientific Research under award FA8655-21-1-7003. R.E.A.G. was supported by the Biotechnology and Biological Sciences Research Council-funded South West Biosciences Doctoral Training Partnership [training grant reference BB/M009122/1]. S.C. acknowledges support from the Program for College Research of the Fonds de recherche du Québec – Nature et technologie (FRQ-NT #275304 and #321076). K.R. was supported by Wellcome Trust Institutional Strategic Support Fund (grant reference AC1910IF14) awarded to D.D.J. The authors would like to thank the Cardiff School of Biosciences Protein Technology Hub for helping with the production and analysis of proteins. The BLIP2, TEM-1 and KPC-2 containing plasmids was a kind gift from Prof Tim Palzkill. The pEVOL plasmid for incorporating azF was a kind gift from Ryan Mehl via Addgene (catalogue #164579).

## Author contributions

REAG, SC and C-SL contributed equally to the research. All authors contributed to the writing of the paper and analysing data. REAG undertook initial rotamer modelling, produced protein and contributed to conductance measurements and analysis. SC undertook replica-exchange molecular dynamics simulations and electrostatic profile modelling. C-SL prepared nanoscale electrodes and contributed to conductance measurements and analysis. KR produced protein. MP conceived and directed the project, and contributed to data analysis. DDJ conceived and directed the project, and contributed to data analysis.

