## Supplementary material for "Optimising CNT-FET Biosensor Design: Predictive Modelling of Biomolecular Electrostatic Gating and its Application to Beta-Lactamase Detection": SI

### These authors contributed equally.

* Corresponding authors

#### Supporting Information.

##### Supporting Methods

***Molecular dynamics modelling of BLIP2 azF mutants***

All simulations were performed in GROMACS [1,2] with the CHARMM36 forcefield [3]. The forcefield was modified to add in the new AzF residue, with partial charges derived from bond charge corrections using ACPYPE [4]. BLIP2 was placed in a dodecahedron box, solvated with TIP3P water and neutralising sodium ions. Energy minimisation was performed with the steepest descent algorithm of maximum 50000 steps and convergence below 1000 kJ $mol^{-1}$ $nm^{-1}$. Long-range electrostatics were determined by Particle-mesh Ewald (cutoff of 1.2 nm) and periodic boundary conditions (PBC) were applied. Temperature and pressure equilibration used the leap-frog algorithm to equilibrate the solvent about the protein. Velocity-rescale temperature coupling and isotropic Berendsen coupling maintained temperature and pressure at 300K and 1 bar using a coupling constant of 100 ps. These parameters were taken forward for the production run using a 2 fs timestep, totalling 200 ns, with energies and coordinates being collected every 10 ps. Simulations were performed in triplicate for each protein.

***Attachment parameterization for H-REMD simulations***

The BLIP2 azF variants were covalently attached to the SWCNT in the biosensor using azido-phenylalanine photochemistry [5]. For the H-REMD simulations, the covalent attachment between BLIP2 and the nanotube is represented with an atomistic resolution to precisely consider the constraints imposed by the attachment site on BLIP2's orientations with respect to the nanotube.

The attachment is modelled by a 4-azido-L-phenylalanine, in which two of the three nitrogen atoms of the azido group were removed (Figure S11). The remaining nitrogen was placed 0.126 nm from the nanotube and the plane of the phenyl group was oriented perpendicularly to the nanotube surface, as is expected[6]. Finally, during the simulations, only the nitrogen atom was kept fixed to maintain attachment with the SWCNT, while all the other atoms of the attachment were free to move.

We parametrized the model compound shown in Figure S11c. A phenylalanine was capped at its N-terminal using an acetyl group and at its C-terminal using a N-methyl amide group, as done when the partial charges of amino acids were parametrized for AMBER. Moreover, a $N(CH_3)_2$ group was added to the CZ carbon to mimic the attachment with the nanotube, without having to consider the whole carbon nanotube, which is computationally demanding when performing the required ab initio calculations to determine the partial charges.

The bonded and Lennard-Jones parameters of the attachment were determined from the Generalized AMBER forcefield (GAFF) [7] using AmberTools20 [8] and ACPYPE [4] to produce GROMACS compatible files. The atom types determined for the model compound are displayed in Figure S11d. As the remaining nitrogen atom of the phenyl azide is frozen during the simulation, we do not explicitly define bonded interactions between the nanotube and the attachment.

The partial charges of the atoms were determined using the RESP protocol as performed to optimize the partial charges of AMBER's amino acids[9]. We considered two backbone structures for the model compound: one representative of a helical state ($\varphi=-60°$ $\varphi=-60°$, $\psi=-40°$ $\psi=-40°$ and $\omega=180°$ $\omega=180°$) and the other of an extended state ($\varphi=217°$ $\varphi=217°$, $\psi=160°$ $\psi=160°$ and $\omega=180°$ $\omega=180°$), as done for phenylalanine in AMBER. First, we determined the electrostatic potential generated by the two conformations using Gaussian16 [10] to perform *ab initio* Hartree-Fock calculations at the 6-31G* level, as done for AMBER. Second, the RESP protocol was applied simultaneously on the two conformations to optimize the partial charge to best represent the electrostatic potential determined using ab initio calculations. RESP follows a two-step procedure: (1) the charges of all atoms are optimized, except those that are constrained, then (2) the charges of the $CH_3$ groups are reoptimized. During RESP, we constrained the partial charges of the N-terminal acetyl group and the C-terminal N-methyl amide group as well as the partial charges of the mainchain C=O and N-H groups to those in AMBER. Moreover, the total charge of the model compound is constrained to zero and chemically equivalent atoms are constrained to the same charge. The optimized charges are displayed in Figure S11d. The electrostatic potential generated by the optimized charges has a relative root mean square of 0.12 with respect to the ab initio electrostatic potential, which is similar to the values obtained for AMBER [9].

##### *H-REMD simulations*

We performed Hamiltonian replica-exchange molecular dynamics (H-REMD) simulations to exhaustively sample the orientations of BLIP2 with respect to the carbon nanotube, while considering the interactions between them. The goal of H-REMD simulations is to enhance the thermodynamics sampling of a molecular system by decreasing the energetic barriers that impair the sampling of traditional MD simulations [11]. To do so, several MD simulations are simultaneously launched in parallel and ordered as followed: the Hamiltonian (energy) in the first MD is unchanged, while it is progressively changed in the other MDs to favour transitions over the energetic barriers. Exchanges of the system between neighbouring MDs is periodically tried using the Metropolis criteria to allow energetically favourable states to diffuse back towards the first MD, which is described by the physically relevant, unscaled energy.

For our system, we realized that BLIP2 hardly samples different orientations with respect to the nanotube when using standard MD simulations. Moreover, we realized that the temperature replica-exchange molecular dynamics (T-REMD) technique usually used to get exhaustive sampling when characterizing biomolecule-SWCNT interactions isn't appropriate for our system because the required high temperatures to unbind the protein from the nanotube aren't preserving the overall structure of the protein.

In our implementation of H-REMD, we selectively scale the attractive part of the Lennard-Jones interactions between BLIP2 and the carbon nanotube. As a result, BLIP2 dissociates more easily from the nanotube at larger scaling, allowing it to thoroughly sample various orientations with respect to the nanotube as the system diffuses back to lower scaling, while preserving the overall structure of BLIP2.

One H-REMD simulation is performed in explicit solvent for each attachment site. The H-REMD simulations were performed using the GROMACS software [2,12] version 2019.6 augmented with the open-source, community-developed PLUMED library version 2.6.2 [13,14] for H-REMD simulations [15]. Each H-REMD simulation consists of 24 MD simulations in parallel with scales ranging from 1.0 to 0 that are applied on the Lennard-Jones attractive interactions between BLIP2 and the nanotube. The systems are solvated and equilibrated as follow: (1) the systems are solvated in explicit water molecules and in 150 mM NaCl to mimic the 1xPBS experimental condition, (2) an additional eleven Na ions are added to neutralize the systems, (3) the systems are minimized first using steepest descent and then conjugate gradient, (4) the temperature is equilibrated at 300 K in a 1-ns simulation during which the position of the heavy atoms of the protein is restrained, and (5) the pressure is equilibrated semi-isotropically, except on the direction along the nanotube axis, at 1 atm using the Berendsen barostat in a 1-ns simulation during which the position of the heavy atoms of the protein is again restrained.

The initial systems for the H-REMD simulations are shown in (Figure S10). To generate the initial states of the scales, we performed a 50-ns MD simulation during which the attractive part of the Lennard-Jones interactions between BLIP2 and the nanotube were removed (scaling of 0) to generate various orientations of BLIP2 with respect to the nanotube, without any contact between BLIP2 and the nanotube. From that simulation, we clustered the trajectory using a cutoff of 1 nm on the carbon alpha atoms of BLIP2 (without aligning the structures since the nanotube is fixed). We used the clusters' centers as initial configurations for the scales to ensure a representativity of the various orientations accessible to BLIP2 when it does not interact with the nanotube.

The interval 500 to 750 ns of the unscaled MD (scale 1.0) was analyzed because the number of orientations remains stable on that interval with 2 main clusters for azF41 and 4 main clusters for azF213 (Figure S4). Moreover, the number of BLIP2's residues within 0.35 nm from the nanotube remains stable on this interval: 9 ± 1 for the azF41 site and 6 ± 1 for the azF213 site. Finally, BLIP2 is structurally stable on that interval in terms of root mean square deviation (RMSD) and secondary structure. The RMSD against BLIP2's crystal structure (PDB: 1JTD) is 0.13 ± 0.01 nm for the azF41 site and 0.10 ± 0.01 nm for the azF213 site. The secondary structure remains near that of the crystal structure, as determined using DSSP. For the azF41 site: 8 ± 1% (9%) for helix, 40 ± 1% (41%) for beta-sheet, 13 ± 2% (14%) for turns, 18 ± 1 (15%) for bend and 21 ± 1% (20%) for coil. For the azF213 site: 8 ± 1% (9%) for helix, 41 ± 1% (41%) for beta-sheet, 14 ± 2% (14%) for turns, 16 ± 1 (15%) for bend and 21 ± 1% (20%) for coil.

##### ***Electrostatic potential simulations***

The APBS software was used to solve the non-linear Poisson-Boltzmann equation [16] to determine the electrostatic potential generated by the protein on the SWCNT. In these simulations, the van der Waals radius and the partial charge of each atom were set to those of AMBER14sb, while the partial charges of the linker atoms were set to 0 to focus on the electrostatic potential generated by the protein. The relative permittivity of the solvent and non-solvent were respectively set to 78.54 and 2. The solvent region was determined using a water molecule radius of 0.140 nm to determine the solvent boundary of the BLIP2/attachment/nanotube system. A monovalent salt concentration of 150 mM was used to mimic the PBS experimental conditions and the radius of the ions was set to 0.2 nm. The electrostatic potential values on the boundary of the simulation box were determined using the simple Debye-Hückel model.

The electrostatic potential was determined using a two-step procedure: the non-linear Poisson-Boltzmann equation was first solved on a large box having 1000 points/nm$^3$ (1 point/Å$^3$) so as to consider the system in its entirety with a large enough space system and boundaries, then the non-linear Poisson-Boltzmann equation was again solved to refine the calculated electrostatic potential values on a region of 2 nm around the nanotube using 8000 points/nm$^3$ (8 points/Å$^3$). In the first step, the size of simulation box was determined from the size of the BLIP2/TEM-1 complex in the simulations multiplied by 1.7 along each direction, yielding a box of 22.5 nm by 22.5 nm by 16.1 nm. The center of the box was set to the average center of the protein in all the sampled configurations. In the second step, the region of 2 nm around the nanotube corresponds to a box of 17.6 nm by 4.80 nm by 4.80 nm that was centred on the nanotube. The length of the nanotube was doubled compared to the H-REMD simulations to consider the fixed Debye-Hückel boundary conditions used in the electrostatic potential calculations.

This two-step procedure was necessary to accurately solve, in a reasonable time the electrostatic potential of the thousands of configurations generated by the H-REMD simulations. The two-step procedure yields similar electrostatic potential values on the nanotube compared to only doing the first step using the greater resolution of the second step (8000 points/nm$^3$). The difference between the two approaches is only 0.01 mV on average, which is small compared to the magnitude of the electrostatic potential on the carbon nanotube (<100 mV).

##### ***Purification of TEM-1 and KPC-2***

Periplasmic extraction is an outer membrane lysis technique used to purify β-lactamase proteins from the periplasmic space, without lysing the whole cell. The cell pellet from inoculated culture was resuspended in 30 mL periplasmic extraction buffer and stirred slowly at room temperature for 10 minutes. The cells were then re-pelleted by (50 mM Tris-HCl, pH 8, 20% (w/v) sucrose1 mM EDTA) centrifugation at 10000 x g at 4°C for 10 minutes, in the Fiberlite™ F21-8 x 50y rotor. Supernatant was discarded, and the cells resuspended in 30 mL chilled magnesium sulphate solution, being stirred slowly over ice for 10 minutes to release the periplasmic proteins. Next, the centrifugation step was repeated again, with the supernatant containing the soluble periplasmic proteins. The supernatant was then mixed in equal volumes with 50 mM Tris-HCl, pH 8 buffer in preparation for column chromatography.

A Resource Q anion exchange column was equilibrated with 30 mL $dH_2O$ and 30 mL 50 mM Tris-HCl, pH 8, buffer. Next, the periplasmic supernatant (mixed 50:50 with water:Tris buffer) was loaded onto the column, with absorbance monitored at 280 nm. Protein was eluted from the column using a gradient of 0 to 1 M NaCl over 30 mL. Fractions containing the desired protein were identified and collected. Size exclusion chromatography was performed using a HiLoad™ 16/600Superdex™ S75 pg column. The column was equilibrated with 50 mM Tris-HCl, pH 8. The BL sample was concentrated in a 10 kDa molecular weight cut off column (Fisher Scientific) until the total protein volume was *circa* 1 mL. The protein sample was loaded with absorbance being monitored at 280 nm. All visible absorbance peaks were fractionated and analysed by SDS-PAGE.

#### Supporting Figures

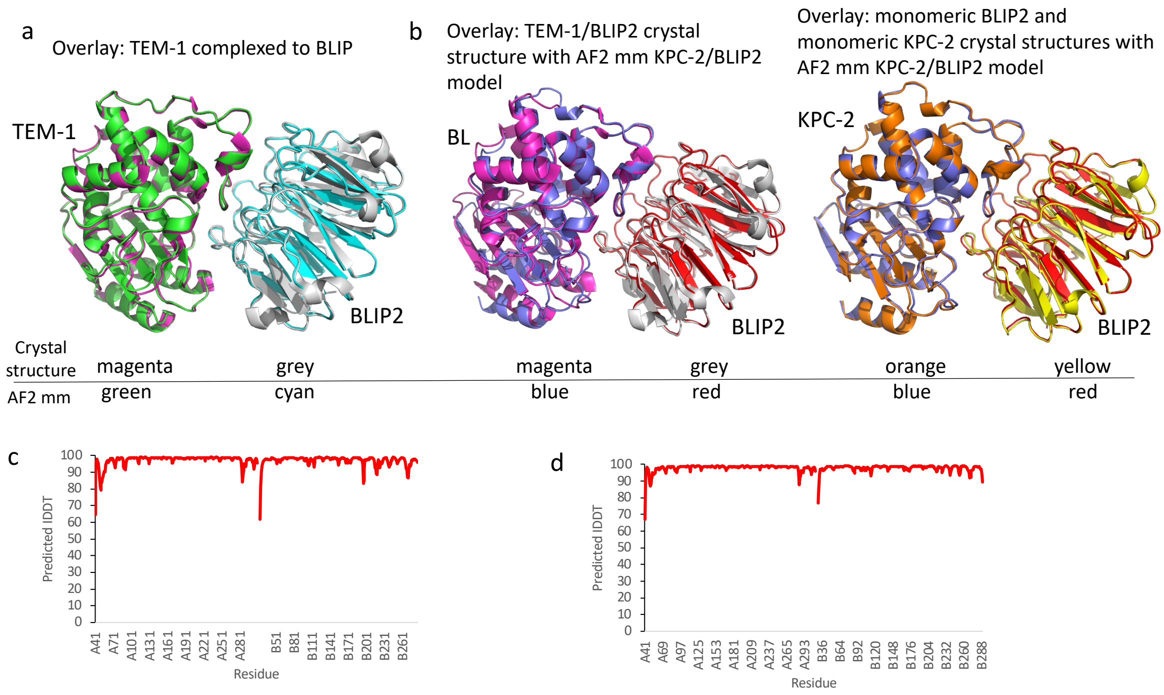


**Figure S1.** AlphaFold2 multimer (AF2mm) modelling of BLIP2 in complex with (a) TEM-1 and (b) KPC-2. Each subunit and its respective colour is outlined on the figure. (a) Overlay of known structure of BLIP2-TEM-1 (1jtd [17]) complex with AF2mm model. The root mean square deviation (RMSD) over Cα between the model and structure is 0.5 Å (b) Overlay of AF2mm model of BLIP2-KPC-2 with known structure of BLIP2-TEM (left; RMSD 0.5 Å) and with crystal structure of BLIP2 alone (3qi0 [18]) and KPC-2 monomer (2ov5 [19]). The predicted local distance difference test (IDDT) for (c) BLIP2-TEM-1 complex and (d) BLIP2-KPC-2 complex. ColabFold v1.5.2: AlphaFold2 using MMseqs2 was used to generate the complex models. (https://colab.research.google.com/github/sokrypton/ColabFold/blob/main/AlphaFold2.ipynb?authuser=0) [20].

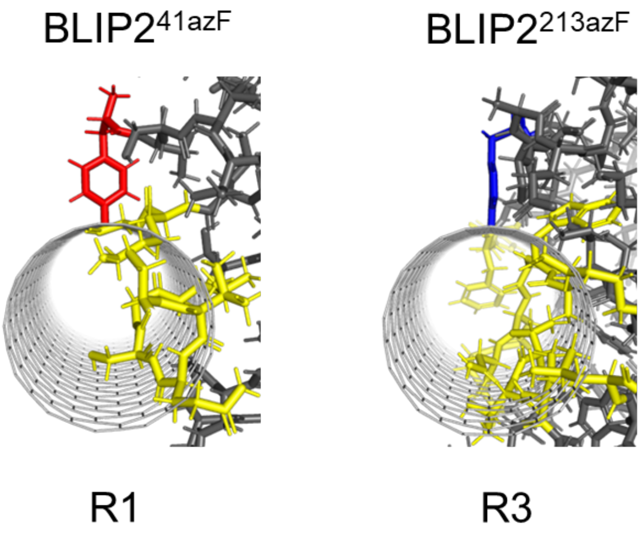


**Figure S2.** Rotamers azF that form steric clashes with an SWCNT on manual docking. The red and blue residues represent rotamer 1 of BLIP2$^{41azF}$ and rotamer 3 of BLIP2$^{213azF}$, respectively, with yellow residues sterically clashing with the nanotube. Docking of BLIP2 azF mutant onto the SWCNT is based on the known nitrene insertion chemistry reaction to provide initial models of the CNT-receptor protein configuration. The nitrene radical formed on exposure to UV light is expected to insert perpendicular to the SWCNT [6].

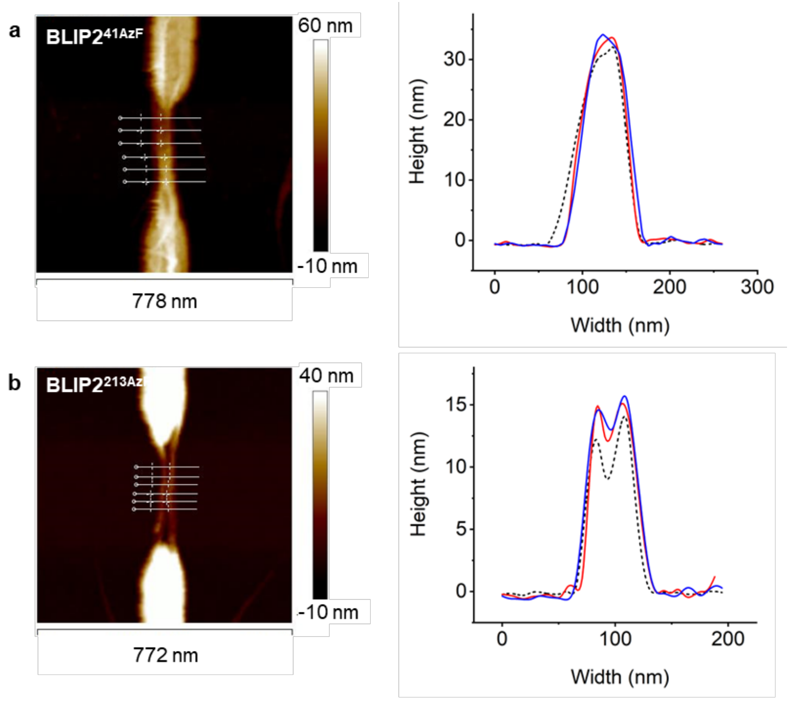


c

| | Change in height (nm) | |
|---|---|---|
| | BLIP2$^{41AzF}$ | BLIP2$^{213AzF}$ |
| BLIP2 | 1.5 nm | 1.9 nm |
| BLIP2 + TEM-1 | 2.1 nm | 2.3 nm |

**Figure S3. AFM analysis of the CNT-FET transistor channel.** AFM images on the left show the functionalised transistor channel, with the overlaid horizontal white lines indicating six cross sections taken for height analysis. An average cross section was calculated for the channel before and after BLIP2 azF functionalisation, and again following β-lactamase sensing, and are plotted in the graphs on the right. Height profiles are seen for SWCNTs (black dashed), BLIP2 azF mutant (red), TEM-1 (blue). (a) BLIP2$^{41azF}$ derivatised CNT-FET, (b) BLIP2$^{213azF}$ derivatised CNT-FET and (c) relative changes in height of the base SWCNT on attachment of BLIP2 and binding of TEM-1. **Supplementary Discussion.** Attachment of BLIP2$^{41azF}$ or BLIP2$^{213azF}$ resulted in an averaged height increase of 1.5 nm or 1.9 nm, respectively. This is lower than the heights anticipated from the dimensions of BLIP2 but is in line with previous AFM measurements that routinely underestimate the heights of soft materials such as proteins [21–23]. After exposure to TEM-1, the height of the CNT-FET increased slightly from 1.5 nm to 2.1 nm and 1.9 nm to 2.3 nm for BLIP2$^{41azF}$ and BLIP2$^{213azF}$, respectively. These relatively small increases (0.4 - 0.6 nm) potentially provides an indication on the position of BLIP2's β-lactamase binding face; with adjacent binding to the SWCNT more likely to fit this data than the β-lactamase binding above the plane of the SWCNT.

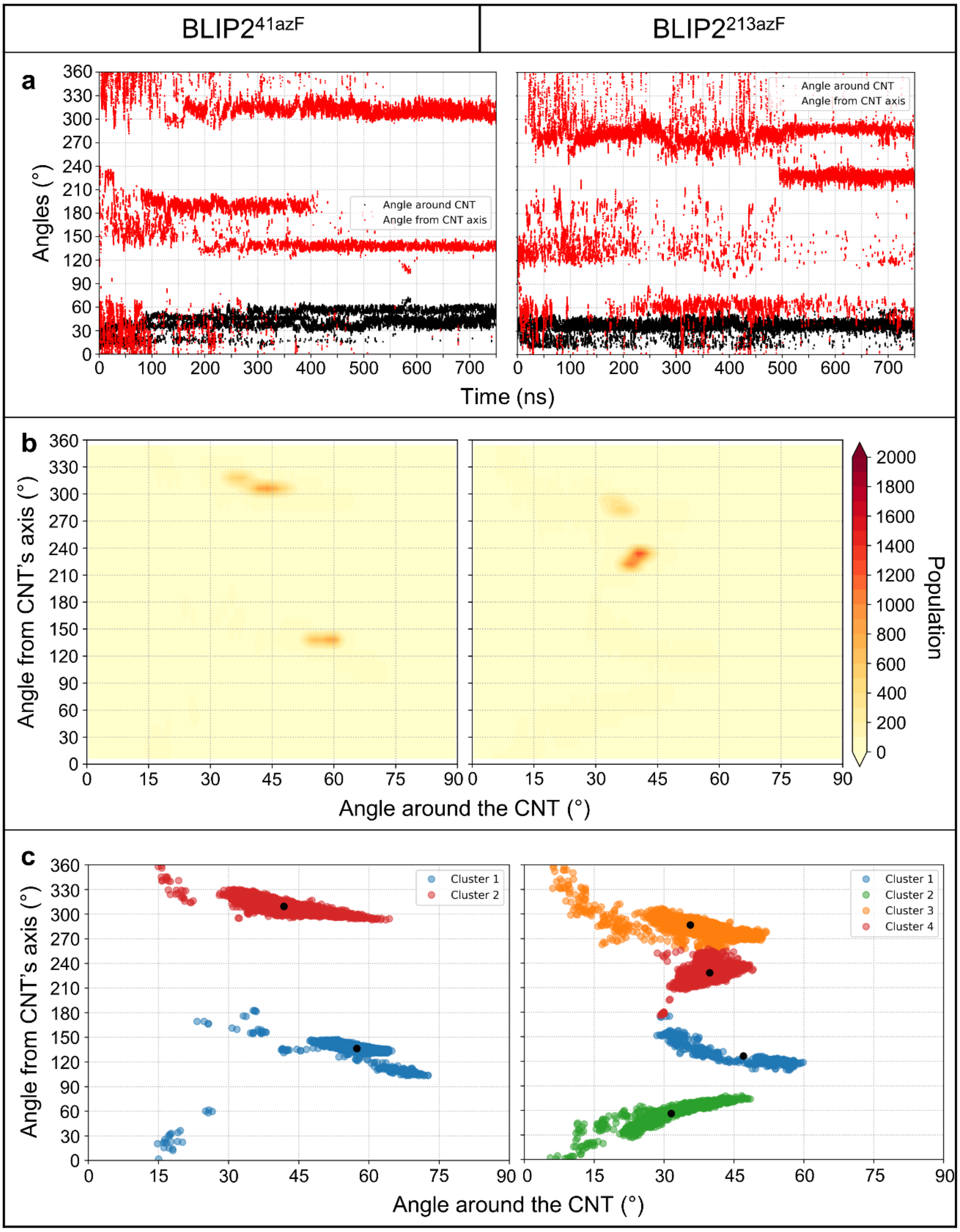


**Figure S4. Orientation of BLIP2 with respect to the nanotube.** For the azF41 and azF213 attachment sites, (A) the orientation of the center-of-mass of BLIP2 in terms of its angle around the nanotube (red) and around the attachment site (black) as a function of time; (B) the orientation histogram; (C) the main orientation clusters determined using k-means.

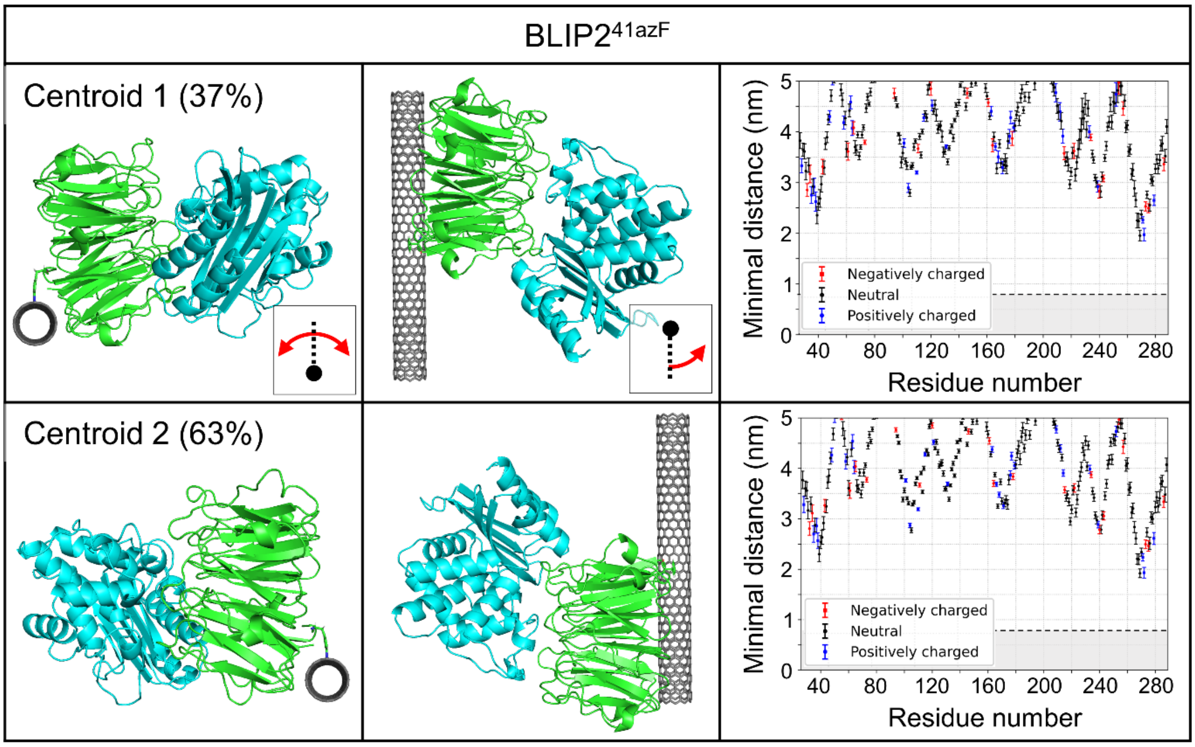


**Figure S5. Main orientations of BLIP2 for the azF41 attachment site.** The centroids correspond to those shown in Figure S4 for BLIP2 (green) on which TEM-1 (teal) has been added. Two different points of view are show for each orientation: from the nanotube axis illustrating the position of BLIP2 around the nanotube, and from the attachment site axis illustrating the position of BLIP2 around the attachment site. Insets show the definition of these angles: the angle around the nanotube runs from -180° (counter clockwise) to +180° (clockwise) with 0° corresponding to BLIP2's center-of-mass being directly above the nanotube (see dotted line), and the angle around the attachment site runs from 0° to 360° (counter clockwise) with 0° corresponding to BLIP2's center-of mass being directly above the nanotube, before the attachment site (see dotted line). The last column shows the minimal distance of each residue of TEM-1 from the nanotube. The shaded region illustrates the Debye length (0.7 nm) in a 1xPBS solution at room temperature.

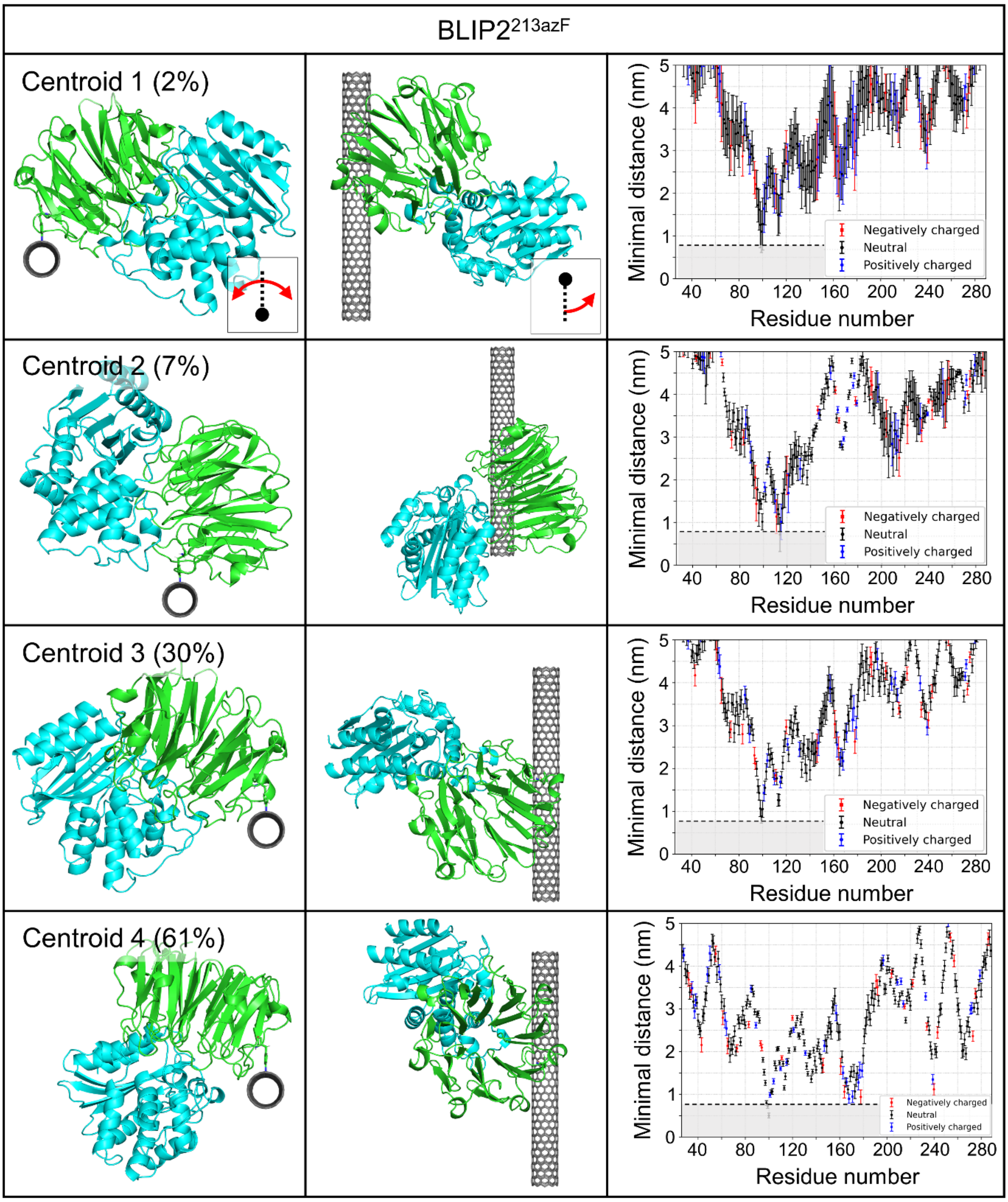


**Figure S6. Main orientations of BLIP2 for the azF213 attachment site.** The centroids correspond to those shown in Figure S4 for BLIP2 (green) on which TEM-1 (teal) has been added. Two different points of view are show for each orientation: from the nanotube axis illustrating the position of BLIP2 around the nanotube, and from the attachment site axis illustrating the position of BLIP2 around the attachment site. Insets show the definition of these angles: the angle around the nanotube runs from -180° (counter clockwise) to +180° (clockwise) with 0° corresponding to BLIP2's center-of-mass being directly above the nanotube (see dotted line), and the angle around the attachment site runs from 0° to 360° (counter clockwise) with 0°

corresponding to BLIP2's center-of mass being directly above the nanotube, before the attachment site (see dotted line). The last column shows the minimal distance of each residue of TEM-1 from the nanotube. The shaded region illustrates the Debye length (0.7 nm) in a 1xPBS solution at room temperature.

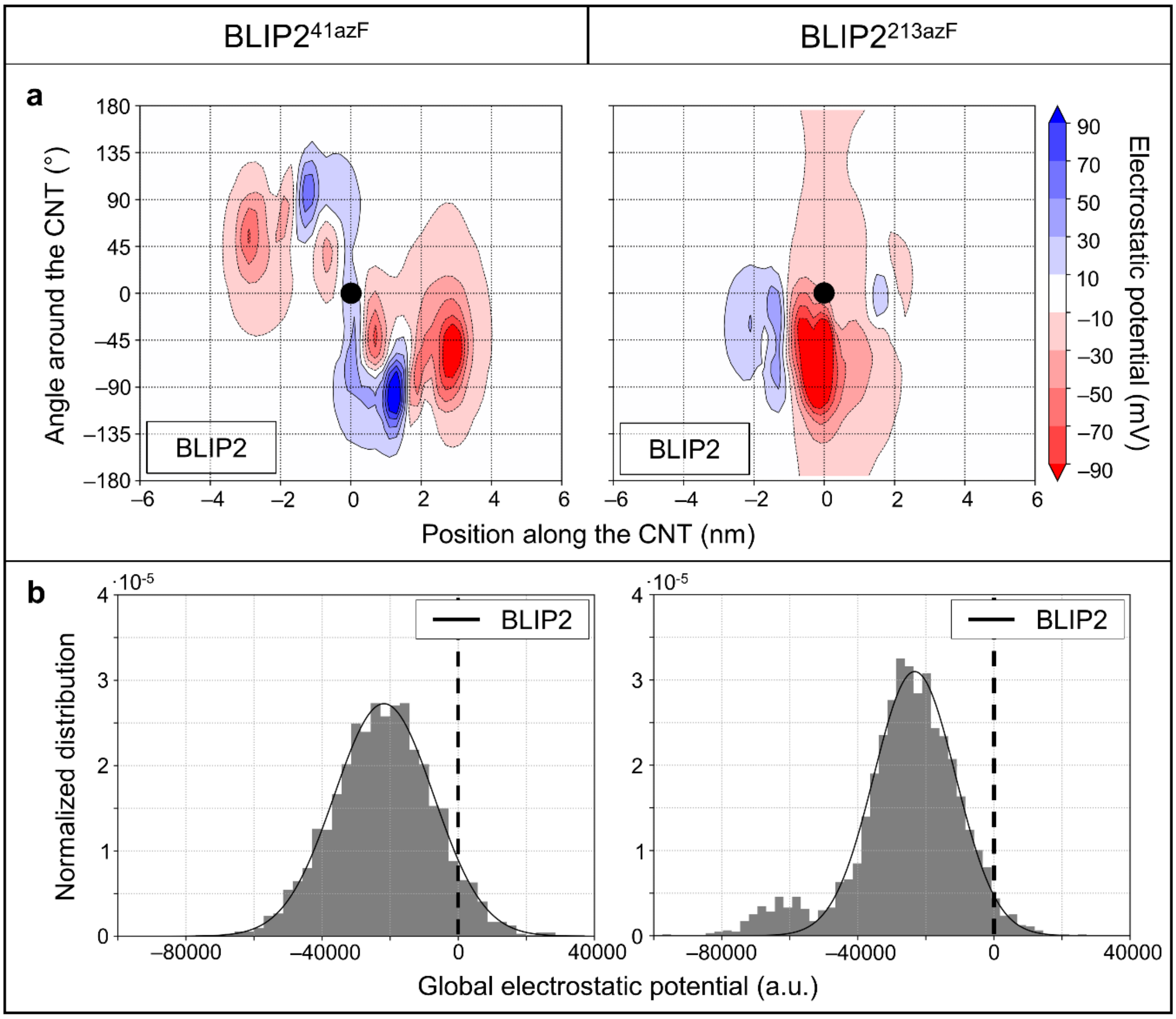


**Figure S7. Electrostatic potential on SWCNT.** Shown is (a) the average electrostatic potential and (b) the global electrostatic potential generated on the SWCNT by BLIP2$^{41azF}$ and BLIP2$^{213azF}$.

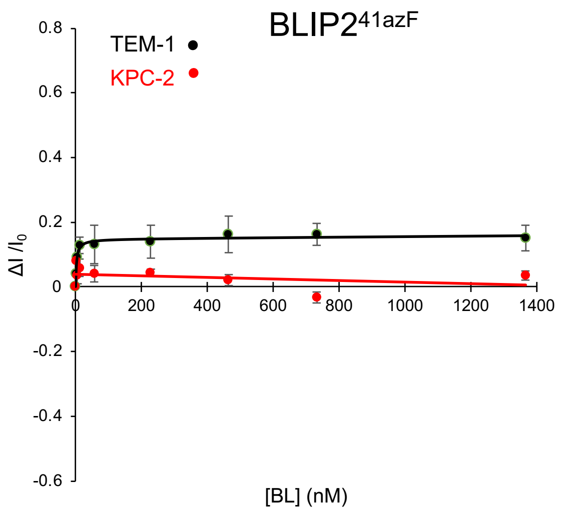


**Figure S8.** BL concentration-dependent change in conductance (at $V_{DS}$ of 0.1 V) for BLIP2$^{41azF}$ CNT-FETs with TEM-1 (black) and KPC-2 (red). The conductance data were collected via I-V measurements between source-drain The data points where fit to one site binding equation in GraphPad Prism. No stable fit to KPC-2 binding was possible. Error bars are standard deviations between multiple independent readings at each concentration. The y axis is scaled to that shown in Figure 4c for comparison.

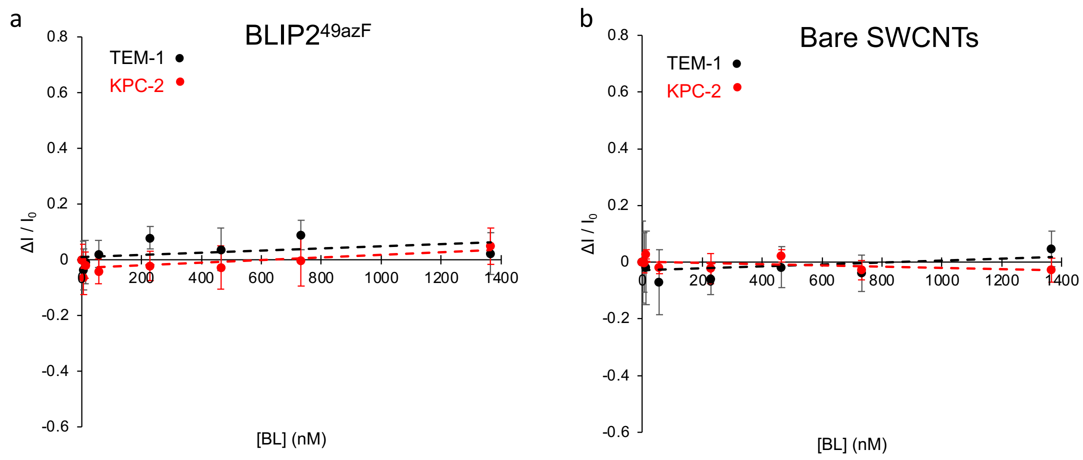


**Figure S9.** BL concentration-dependent change in conductance (at $V_{DS}$ of 0.1 V) for (a) BLIP2$^{49azF}$ and (b) bare SWCNT-FETs with TEM-1 (black) and KPC-2 (red). BLIP2$^{49azF}$ contains the azF mutation within the binding interface and thus will block BL binding on attachment to a SWCNT. The conductance data were collected via I-V measurements between source-drain. Error bars are standard deviations between multiple independent readings at each concentration. No stable fit to the data was possible so trendlines are shown as dashed lines. The y axis is scaled to that shown in Figure 6c for comparison.

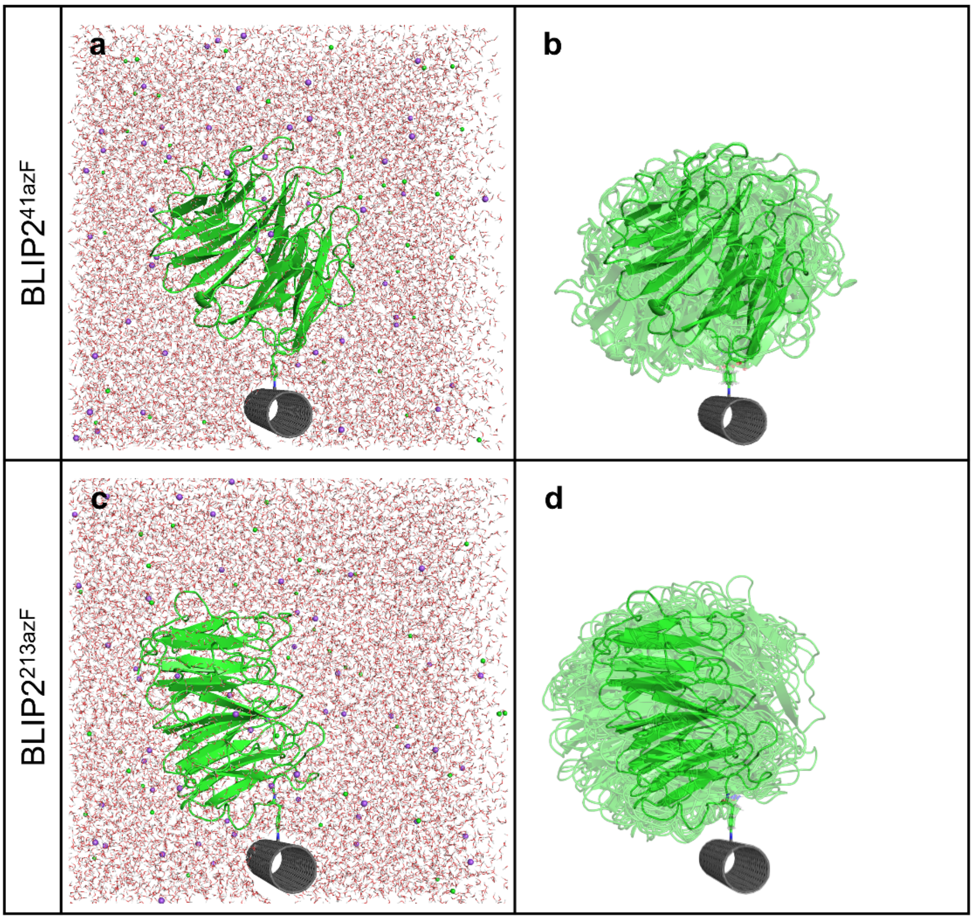


**Figure S10. Simulated systems.** (a-c) The simulated systems containing the BLIP2 protein covalently attached to the carbon nanotube, and explicitly solvated in water molecules and 150 mM NaCl. (b-d) The initial structures used for all scales.

a

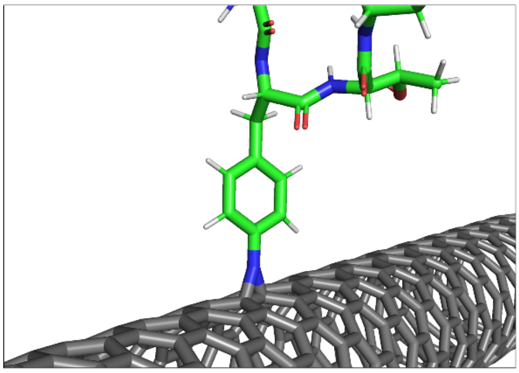

b

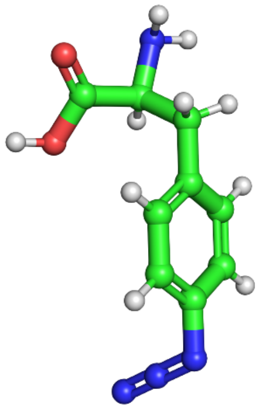

c

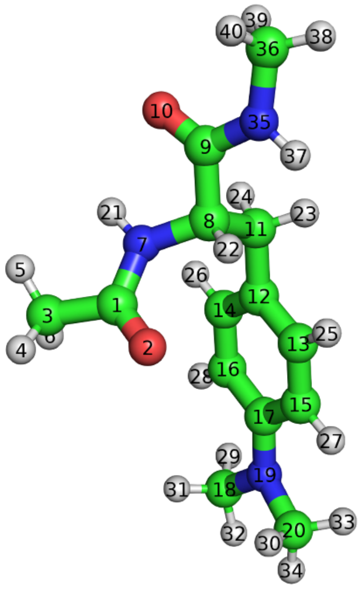


d

| Atom no | Atom type | Charge (e) | Atom no | Atom type | Charge (e) |
|---|---|---|---|---|---|
| 1 | c | 0.5972 | 19 | nh | -0.3486 |
| 2 | o | -0.5679 | 20 | c3 | -0.0152 |
| 3 | c3 | -0.3662 | 21 | hc | 0.2719 |
| 4-5-6 | hc | 0.1123 | 22 | h1 | 0.1212 |
| 7 | n | -0.4157 | 23-24 | hc | 0.0272 |
| 8 | c3 | -0.0274 | 25-26 | ha | 0.1440 |
| 9 | c | 0.5973 | 27-28 | ha | 0.1440 |
| 10 | o | -0.5679 | 29-31-32 | h1 | 0.0569 |
| 11 | c3 | -0.0541 | 30-33-34 | h1 | 0.0569 |
| 12 | ca | 0.0481 | 35 | n | -0.4157 |
| 13-14 | ca | -0.1969 | 36 | c3 | -0.1490 |
| 15-16 | ca | -0.1539 | 37 | hn | 0.2719 |
| 17 | ca | 0.1861 | 38-39-40 | h1 | 0.0976 |
| 18 | c3 | -0.0152 | | | |

**Figure S11. Attachment parametrization.** (a) The covalent attachment between the carbon nanotube and BLIP2. (b) The 4-azido-L-phenylalanine molecule. (c) The compound used to parameterize the attachment. (d) The atom type from GAFF and the RESP optimized charge for each atom. Atoms on a same line are equivalent and were constrained to the same charge. The values in red were constrained.
